# Initial virome characterization of the common cnidarian lab model *Nematostella vectensis*

**DOI:** 10.1101/2020.01.14.906370

**Authors:** Magda Lewandowska, Yael Hazan, Yehu Moran

## Abstract

The role of viruses in forming a stable holobiont has been a subject of extensive research in the recent years. However, many emerging model organisms still lack any data on the composition of the associated viral communities. Here, we re-analyzed seven publicly available transcriptome datasets of the starlet sea anemone *Nematostella vectensis*, the most commonly used anthozoan lab model, and searched for viral sequences. We applied a straightforward, yet powerful approach of *de novo* assembly followed by homology-based virus identification and a multi-step, thorough taxonomic validation. The comparison of different lab populations of *N. vectensis* revealed the existence of the core virome composed of 21 viral sequences, present in all adult datasets. Unexpectedly, we observed almost complete lack of viruses in the samples from the early developmental stages which together with the identification of the viruses shared with the major source of the food in the lab, the brine shrimp *Artemia salina*, shed new light on the course of viral species acquisition in *N. vectensis*. Our study provides an initial, yet comprehensive insight into *N. vectensis* virome and sets the first foundation for functional studies of viruses and antiviral systems in this lab model cnidarian.

## 1. Introduction

Viruses, the absolute parasites of virtually all living organisms, constitute the most abundant and diverse entity on Earth [1, 2]. Owing to their dependency on host organisms and the resulting continuous evolutionary arms race with their hosts, viruses are predominantly studied in the context of a pathogenic state of the host cells [3]. Moreover, much focus is given to deciphering viruses of humans and other economically important species while the viral diversity of other hosts remains understudied [4]. Importantly, many viruses can remain in a dormant state, which is neutral to the cell environment, or maintain a commensal relationship with the host [5–7]. In 1990 Lynn Margulis first introduced a concept of ‘holobiont’ which referred to a metaorganism formed by a symbiosis of separate living entities which constitutes an individual unit of selection [8]. Since the recognition that the collective of prokaryotic and eukaryotic viral species present in the host, hereinafter called ‘virome’, also form a part of the metaorganism, its complex role in host disease, development, and evolution is a subject of studies and debates [9].

*Nematostella vectensis*, also known as the starlet sea anemone, is a non-symbiotic emerging cnidarian model species owing to the facility of its culture under laboratory conditions and a wide range of accessible tools for genetic engineering (reviewed in [10]). Cnidaria is a phylum representing early-branching Metazoa and has diverged from its sister group Bilateria, which includes the vast majority of extant animals, approximately 600 million years ago [11], making it an attractive group for wide range of comparative studies. Cnidarians are divided to two major classes: Anthozoa (sea anemones and corals) and Medusozoa (jellyfish and hydroids) [12]. Among cnidarians, much focus has been placed on deciphering the composition and significance of virome of corals (reviewed in [13]), their photosynthetic dinoflagellate symbionts from the *Symbiodinium* genus [14, 15], and their symbiotic sea anemone proxy *Exaiptasia pallida* (formerly called *Aiptasia pallida*) [16], mainly due to the environmental significance of the coral reef ecosystems. Members of coral virome have been suggested to play a role in some coral diseases [17, 18], and a general increase of viral abundance has been observed during bleaching (loss of dinoflagellate symbionts) of several coral species [19–21].

*Nematostella* is arguably the most commonly used anthozoan lab model [10] and extensive research has been done so far to uncover its microbiome composition, significance and dynamics of the interplay with the host environment [22–24]. In contrast to microbial studies and despite its wide use as a lab model, the composition of stable virus community forming the holobiont of this sea anemone has not yet been studied. Likewise, no virus capable of infecting *Nematostella* has been identified so far, impairing our understanding of viral pathogenesis in the starlet sea anemone. Furthermore, lack of any insights to *Nematostella* viral community hinders the research on the role of RNA interference (RNAi) – the major sequence-specific antiviral system of plants and invertebrates [25–27] – in the innate immune response of *Nematostella* to viruses.

In this study, we aimed to characterize for the first time *N. vectensis* virus communities. To this end, we re-analyzed various publicly available RNA-seq datasets and applied a strategy of *de novo* assembly of putative viral sequences followed by thorough phylogenetic validation. Our approach was expanded by generating novel transcriptome datasets from the primary food source of laboratory sea anemones, the brine shrimp *Artemia salina*. We have identified a set of unique *Nematostella*-specific and *Artemia*-specific viral sequences with sound homology to known viruses, as well as characterized the core *N. vectensis* virome present in all previously sequenced lab populations. Finally, we observed a lack of viral load in early developmental stages and determined approach-dependent differences between individual datasets which might serve as a guide for future RNA virome research in this species.

## 2. Materials and Methods

In this study, we used two types of datasets: publicly available RNA-seq paired-end data from *N. vectensis* spanning different developmental stages and two novel RNA-seq datasets from *A. salina* nauplii, which constitutes the main food source for *N. vectensis*, generated in-house.

### 2.1. RNA-extraction and sequencing

Approximately 100 μl of *A. salina* nauplii were used for each of the two biological replicates. Total RNA was extracted with Tri-Reagent (Sigma-Aldrich, St. Louis, MO, USA) according to manufacturer’s protocol, treated with 2 μl of Turbo DNAse (Thermo Fisher Scientific, Waltham, MA, USA) and re-extracted with Tri-Reagent. The quality of total RNA was assessed on Bioanalyzer Nanochip (Agilent, Santa Clara, CA, USA), although no RNA Integrity Number (RIN) was available due to the presence of a single peak representing 18S rRNA subunit, commonly observed in arthropods [28]. RNA-seq libraries were constructed using SENSE Total RNA-seq Library Prep Kit v2 (Lexogen, Vienna, Austria) following the manufacturer’s protocol and sequenced on NextSeq 500 (Illumina, San Diego, CA, USA) with 75 nt read length. The raw data have been deposited at NCBI SRA database (accession numbers are not yet available and will be provided during the review).

### 2.2. N. vectensis transcriptome datasets

We used previously published RNA-seq datasets of 50 – 100 nt paired-end reads from six different studies. The first dataset was reported by Babonis *et al.* 2016 [29] (NCBI BioProject accession: PRJEB13676) and includes three transcriptome replicates of nematosomes, mesenteries, and tentacles of adult *Nematostella*. The second study by Tulin *et al.* 2013 spans the first 24 h of embryogenesis and the dataset is deposited on the Woods Hole Open Access Server (http://darchive.mblwhoilibrary.org/handle/1912/5613) [30]. Next dataset, by Oren *et al.* 2013 (NCBI BioProject accession: PRJNA246707), reports circadian rhythm transcriptome of adult sea anemones [31]. Two datasets generated at Vienna University, partially published by Schwaiger *et al.* 2014 (NCBI BioProject accession: PRJNA200689 and PRJNA213177) were depleted from duplicates and merged [32]. Samples spanning all *Nematostella* developmental stages were further divided into two groups encompassing polyA-selected and rRNA-depleted samples. The fifth transcriptomic adult sample came from the study by Fidler *et al.* 2014 (NCBI BioProject accession: PRJNA200318) [33]. The last dataset was reported by Warner *et al.* 2018 and includes samples spanning the first 144 h of regeneration of 6-week-old juveniles (NCBI BioProject accession: PRJNA419631) [34]. Detailed list of NCBI accession numbers, raw and filtered read counts of all samples used in this study is shown in Supplementary file 1, Table S1.

### 2.3. Raw reads processing and filtering

Quality of raw reads was assessed by FastQC software [35]. Reads were trimmed and quality filtered by Trimmomatic with the following parameters (LEADING:5 TRAILING:5 SLIDINGWINDOW:4:20 MINLEN:36) [36]. Only paired-end reads were used for downstream analysis. Bowtie2 [37] with following parameters (--local -D 20 -R 3 -L 10 -N 1 -p 8 --mp 4) was used to align the reads to *Nematostella vectensis* genome (NCBI accession: GCA_000209225.1) [38], as well as to all sequences of transfer, mitochondrial and cytoplasmic ribosomal RNA retrieved from RNAcentral database [39], retaining unmapped reads after each step. The same mapping strategy was used to remove external RNA Controls Consortium (ERCC) spike-ins whenever used during library construction. In order to further filter retained datasets of all *Nematostella*-derived short reads, we performed a local stringent BLASTn (version 2.3.0+) search against *Nematostella* genome. Next, we removed from the datasets all reads below the e-value cutoff of 1e-15 by a custom Python script (script available at https://github.com/yael-hzn/Nematostella-viruses-project/blob/master/NV_viruses_filter_reads_out_script.ipynb). Of note, at these stages we removed all possible viral sequences that might have been falsely assembled or incorporated into the official version of the *Nematostella* genome [38].

### 2.4. Sequence assembly and viral sequence identification

The remaining reads of each dataset were *de novo* assembled by Trinity (version trinityrnaseq_r20140717) with default parameters [40]. The assembly was repeated inputting merged filtered reads from all datasets. This assembly was processed equally to other datasets and treated henceforth as a general *N. vectensis* viral dataset. Next, we employed a thorough three-step BLAST-based filtering process in order to retrieve only the sequences of high, certain virus homology. After the assembly, we ran simultaneous local BLAST searches – first, BLASTn against *Nematostella* genome to classify sequences which previously were too short for generating a significant homology score and two BLASTx searches against viral protein database (RefSeq release 86, Swiss-Prot Release 2018_02) and prokaryote protein database (Swiss-Prot Release 2018_02) with e-value 1e-5 and 1e-10 as a cutoff, respectively. Only contigs longer than 200 nt and of unambiguous viral origin were retained for downstream analysis and were trimmed to recover only sequence with known homology. Custom Python script for comparison of multiple BLAST searches and trimming of putative virus-derived contigs is available at https://github.com/yael-hzn/Nematostella-viruses-project/blob/master/Nv_viruses_alignment_summary_script.ipynb.

The second step of virus identification was a local BLASTx search against all proteins available in SwissProt database (Release 2018_02) in order to remove sequences of clear non-viral origin. After the search with e-value 1e-10 as a cutoff, we retrieved all putative viral sequences, as well as contigs without any clear homology to any protein in the database. As a final filtering step, we performed a remote BLASTx 2.7.1+ search against RefSeq Protein database (Release 89, e-value cutoff was 1e-10) to select only those sequences with identifiable homology to the known eukaryotic viruses. It is worth noting that this step of sequence filtering overlooks novel species without any clear homology to previously annotated viruses.

### 2.5. Taxonomic annotation

The retrieved viral sequences were taxonomically annotated following the approach of Goodacre *et al.* [41]. In brief, taxonomic identifiers obtained during the final BLASTx search were used for climbing the taxonomic tree by using NCBI parent-child taxonomic identifier definitions (file available at ftp://ftp.ncbi.nih.gov/pub/taxonomy/taxcat.zip; nodes.dmp file) until the family level was reached. Family names were recovered by mapping family taxonomic identifier to the taxonomic name (file available at ftp://ftp.ncbi.nih.gov/pub/taxonomy/taxcat.zip; names.dmp file). Taxonomic annotation was manually checked to be consistent with the International Committee on Taxonomy of Viruses (ICTV) Master Species List 2018a v1.

### 2.6. Reads quantification

Filtered reads from each dataset (after non-coding RNA/spike-ins removal) and *Artemia* transcriptome were remapped to the general *N. vectensis* viral dataset by Bowtie2, applying the following parameters (-N 1 -L 15). To run a statistical comparison of the datasets, we downloaded two single-end replicates from BioProject PRJNA419631 [34] and performed the same remapping. Next, we established the core virome by selecting viral sequences from the general assembly, to which reads from all datasets were mapped (excluding reads from embryogenesis dataset due to very low level of viral load [30]). Venn diagram was generated with the online tool jvenn [42]. Detailed list of NCBI accession numbers of the single-end datasets is presented in Supplementary file 1, Table S1.

### 2.7. Validation of candidate viruses

To confirm the presence of viral sequences identified in RNA-seq libraries, we selected 10 contigs from the *Nematostella* core virome for which we performed reverse transcription polymerase chain reaction (RT-PCR) assays. RNA was extracted from adult female sea anemone and 2-days-old planula following the same protocol used for Artemia RNA extraction. cDNA was constructed using SuperScript III (Thermo Fisher Scientific) according to the manufacturer’s protocol. cDNA was amplified with Q5® High-Fidelity DNA Polymerase (New England Biolabs, Ipswich, MA, USA) in 25 μl reaction with thermocycling conditions as follows: 98°C for 30 sec, followed by 35 cycles of 98°C for 10 s, 60°C for 20 sec, 72°C for 20 sec and final extension at 72°C for 2 mins. Fragment of the *Nematostella* NVE5273 gene was amplified under the same conditions as a positive control. PCR products were analyzed on 1.5% agarose gel. Sequences of primers and length of amplified fragments are shown in Supplementary file 1, Table S2.

### 2.8. Statistical analysis

To test whether datasets have significantly different viral composition, we compared normalized remapping results between PRJEB13676 and PRJNA419631 (Babonis *et al.*, Warner *et al.*, respectively), for which biological triplicates were available. PCA factor analysis (max no. of factors = 5) revealed two factors with eigenvalue > 2, which explained together 90.66% of the observed variance (Supplementary file 2). After extracting component loadings for these factors, we ran a separate t-test for each factor. All the calculations were done in SYSTAT version 13.2. Mean and standard deviation (SD) from mean were calculated in RStudio [43].

## 3. Results

A total number of 1,908,174,590 reads pairs distributed over 7 publicly available paired-end read RNA-seq datasets and 54 samples were used for *de novo* assembly and homology-based viral sequences search (Table 1). We identified 94 unique viral sequences in the merged dataset which served as a viral database for all further analyses, while the number of unique viral sequences in individual datasets varied from 6 to 76 (Table 1). Viral contigs in the general viral assemblage ranged in length from 200 to 7,731 nt. All sequences from general viral assembly and individual dataset assemblies are presented in Supplementary file 1, Tables S3, S6-S12, Text files S1, S3, S5, S7, S9, S11, S13, S15. Furthermore, we retained all contigs of viral origin which were not trimmed to contain only fragments homologous to known viruses i.e. sequences with stretches of both viral and unknown homology (Supplementary file 1, Text files S2, S4, S6, S8, S10, S12, S14, S16).

**Table 1.** Summary of sequence data used for virus identification in RNA-seq studies. Retained reads refer to read counts after quality filtering, trimming, and removal of *Nematostella* genome, transfer, mitochondrial and cytoplasmic rRNA, as well as ERCC spike-ins. Identified final viral reads were filtered from the total number of *de novo* assembled contigs and cleared from duplicates.

| Reference | No. of samples | Raw read pairs | Retained reads | Total number of contigs | Identified viral sequences |
| --- | --- | --- | --- | --- | --- |
| Babonis *et al.* 2016 | 9 | 364,726,242 | 18,134,716 | 99,241 | 76 |
| Tulin *et al.* 2013 | 6 | 112,159,243 | 5,251,525 | 7,631 | 6 |
| Oren *et al.* 2015 | 13 | 105,403,849 | 6,805,643 | 10,968 | 12 |
| Shwaiger *et al.* 2014 *polyA* | 8 | 532,867,635 | 75,383,380 | 40,225 | 50 |
| Shwaiger *et al.* 2014 *rRNA-depleted* | 2 | 155,212,236 | 23,904,647 | 6,470 | 62 |
| Fidler *et al.* 2014 | 1 | 95,331,053 | 6,988,326 | 9,902 | 25 |
| Warner *et al.* 2018 | 15 | 542,474,332 | 35,925,820 | 24,934 | 73 |
| All datasets merged | 54 | 1,908,174,590 | 172,394,057 | 155,322 | 94 |

Analysis of the dataset generated by Tulin *et al.*, which captures the stage of *Nematostella* embryonic development, revealed only six short unique sequences with homology to Rous sarcoma virus, a representative of the *Retroviridae* family. Similar scarcity of viral sequences in early developmental stages was also observed in individual samples assemblage (unfertilized egg, blastula and gastrula samples from polyA-selected and rRNA-depleted libraries from Schwaiger *et al.*, data not presented). Therefore, when searching for the common *Nematostella* virus community we decided to exclude this dataset and focus on viromes from the non-embryonic developmental stages.

### 3.1. Viral community classification

Our homology-based identification of viral sequences in *N. vectensis* RNA-seq datasets revealed sequences belonging to 11 viral families, 2 unassigned orders and 3 unclassified groups (Fig. 1). Detected viral families included *Baculoviridae*, *Iridoviridae*, *Marseilleviridae*, *Mimiviridae*, *Phycodnaviridae*, *Pithoviridae*, *Reoviridae*, *Retroviridae*, *Rhabdoviridae* and *Yueviridae*. The most prevalent viral family was dsDNA *Iridoviridae* (23.26%) which was present in 5 out of 6 non-embryonic transcriptomes (Fig. 1c). However, the highest abundance of viral sequences falls into a group of unclassified RNA viruses (41.86%, Fig 1a) which is entirely composed of a group of novel viruses captured in a wide range of invertebrate species by Shi *et al.* (denoted in our results as “unclassified RNA viruses ShiM-2016”) [4]. Similarly, this is also the most abundant group when analyzing the genomic composition of detected viruses (41.86%), with almost equal contribution of dsDNA viruses (37.21%, Fig 1c).

**Figure 1.**
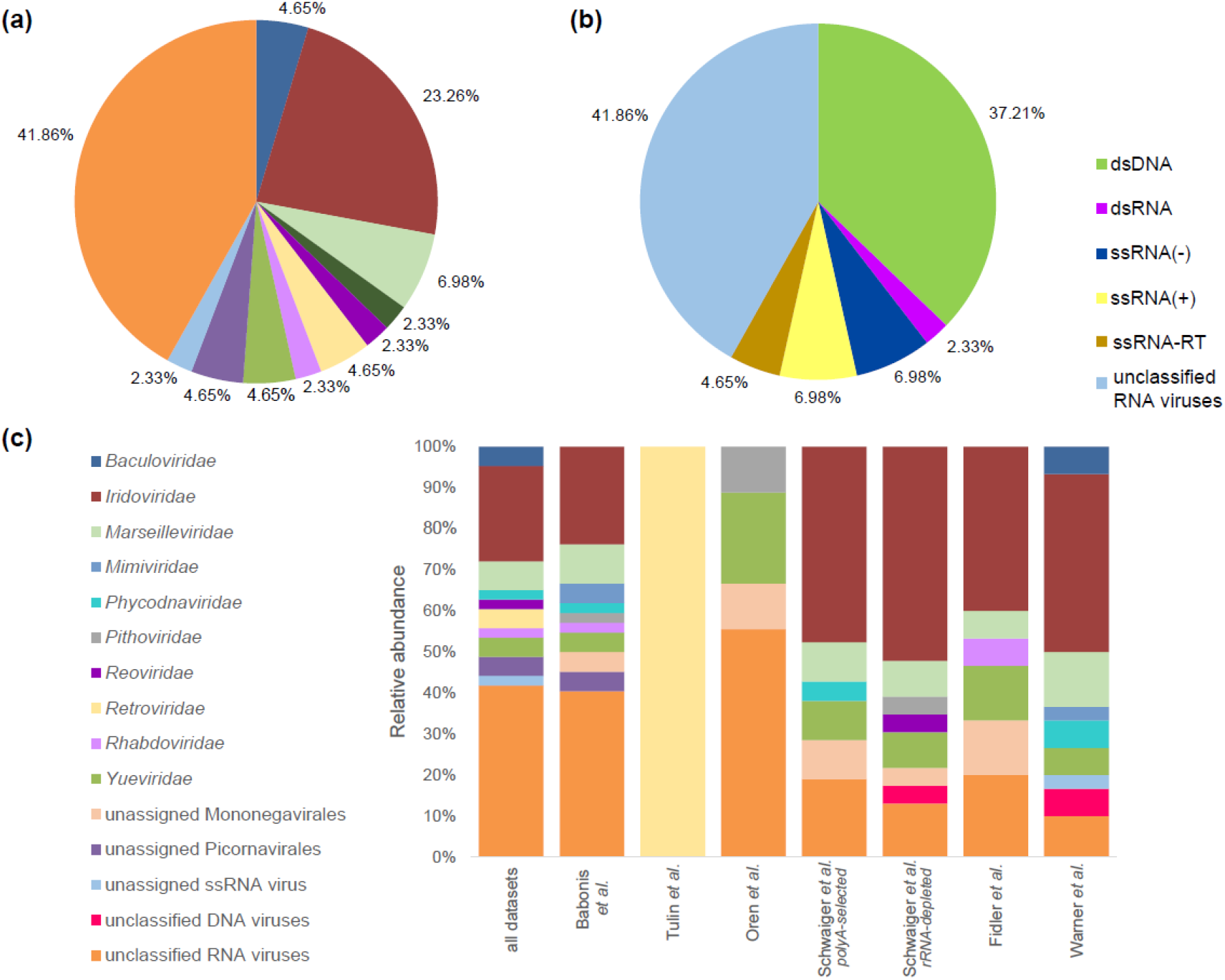
Taxonomic classification of *N. vectensis* viral sequences. Distribution of viral families (a) and groups (b) in the sequence assembly from all merged datasets and (c) relative abundance of viral families within each studied dataset. Sequences representing fragments of the same viral species were collapsed into one entry.

Composition of viral communities was relatively stable across all adult-associated datasets, although several population-specific viruses were detected (Fig. 1c). For instance, Changjiang picorna-like virus 1 was found uniquely in the dataset from Oren *et al.* focusing on the circadian rhythm transcriptome of adult sea anemones. It is the most prevalent virus in the *de novo* assemblage and accounted for 93.03% of all viral reads in this lab population. Such high contribution of one virus to a virome might also partially explain the lack in this dataset of a representative of the most common family *Iridoviridae*, as a result of insufficient sequencing depth and underrepresentation of reads coming from less abundant viruses. Similarly, the sequence of Beihai picorna-like virus 57 was found only in the dataset from Babonis *et al.*, which reported the transcriptome of nematosomes, mesenteries and tentacles of adult *Nematostella*, and comprised 52.5% of all viral reads in this lab population.

To discern *Nematostella*-specific viruses from those derived from the primary source of food, *A. salina* nauplii, we generated two replicates of *Artemia* RNA-seq libraries (21,787,250 and 37,055,827 raw single-end reads). Raw reads were quality-filtered, trimmed and directly mapped to the constructed viral database composed of 94 viral contigs. Mapping to *N. vectensis* virome instead of *de novo* assembly of *A. salina* viruses was motivated by finding an overlap between the sea anemone and its food source at the lab, rather than revealing complete *A. salina* virome. Mapping to our general viral database identified 7 contigs which were shared between *N. vectensis* and *A. salina* (found in both RNA-seq replicates). Those viral sequences ranged in length from 279 to 5,952 nt and represented family *Yueviridae*, *Rhabdoviridae* and unclassified RNA viruses (Supplementary file 1, Table S4).

### 3.2. N. vectensis core virome

In order to establish a core virome of *N. vectensis* i.e., a collective of viral species present in all, but embryonic studied datasets, we decided to use the data from the remapping stage, rather than assembled contigs. We assumed that all viral fragments detected in a sample are a more reliable representation of the true virome, since such approach takes into account lowly expressed or incomplete viral sequences, which might not be assembled into contigs or may be partially undetected when sequencing depth is insufficient. The obtained set is composed of 21 viral sequences (Fig. 2, Supplementary file 1, Table S5), 6 of which are *A. salina*-derived viruses. 61.9% of the sequences from the core virome represent *Iridoviridae*, the family of dsDNA viruses, spanning two genera – *Lymphocystivirus* and *Iridovirus*. Among all *Iridoviridae* sequences, none of them was mapped in *Artemia* libraries. This places *Iridoviridae* as the most common viral family, specific to *N. vectensis* rather than derived from the food source. Interestingly, the highest contribution of the population-specific virus to a total detected population virome was observed in more noise-prone rRNA-depleted libraries (Schwaiger *et al.* [32]) and in the tissue-specific dataset (Babonis *et al.* [29]). We further validated the presence of 10 randomly chosen sequences from the core virome set in cDNA preparations from *A. salina*, adult female sea anemone and 2-days-old planula originating from our lab population. The RT-PCR analysis confirmed complete lack of viral load in the planula stage and the presence of *N. vectesis*-specific and *A. salina*-derived viruses in our samples (Figure S1).

**Figure 2.**
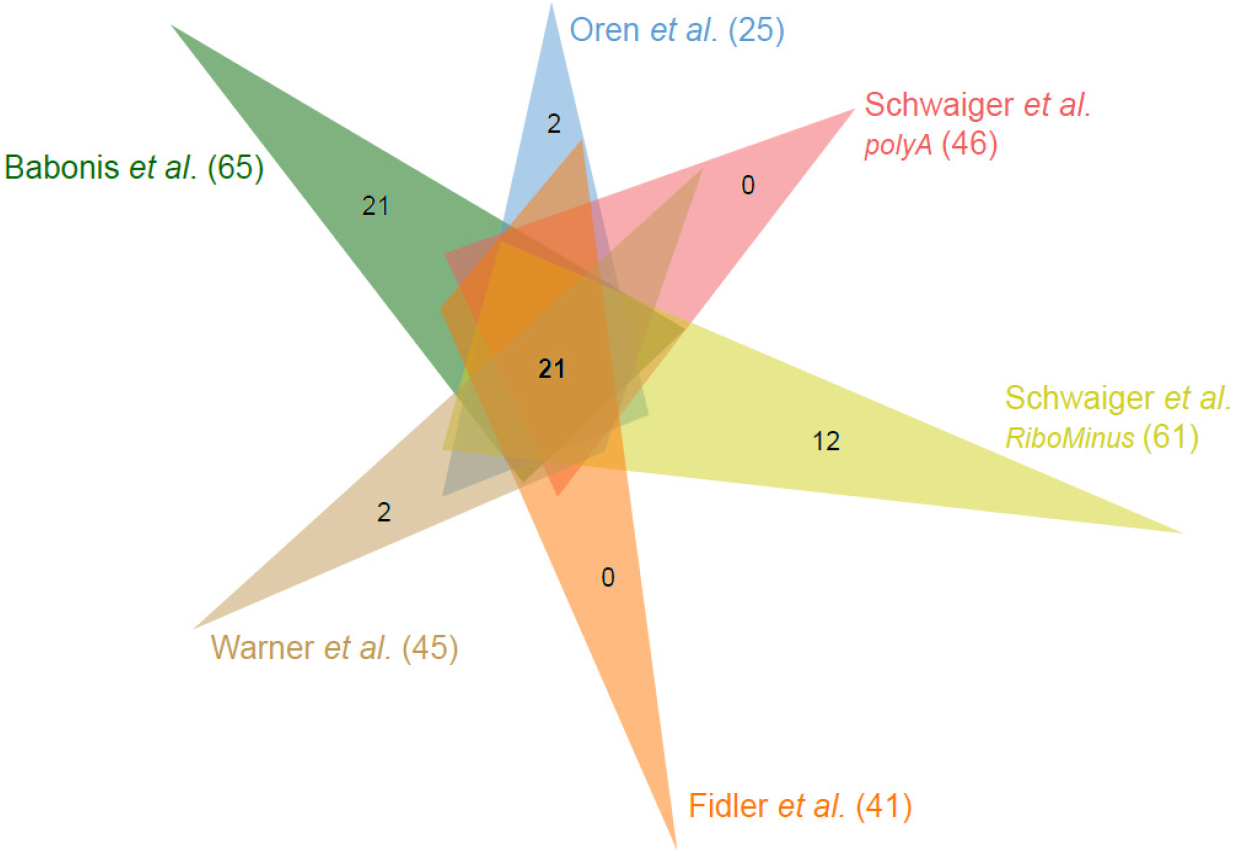
Venn diagram of core viral sequences for all non-embryonic datasets revealed by remapping filtered reads to the viral contigs assembled from the merged dataset. For simplicity, only numbers of contigs common to all datasets and specific to each dataset are shown. Values in brackets indicate the total number of different viral contigs to which reads from each dataset were successfully remapped.

### 3.3. Interpopulation comparison

In order to characterize the general pattern of inter-population differences in the viral load, we compared the normalized counts of viruses-mapped reads of each dataset. As expected, we observed a prominent enrichment in viral sequences in the rRNA-depleted libraries (SDs from mean = 2.218, Fig. 3a, Table 2) when compared to the rest of polyA-selected libraries. Next, we compared the contribution of viruses derived from the food source to the total viral load of *Nematostella*. Similarly to the general viral load pattern, rRNA-depleted libraries displayed a significantly higher load of *A. salina*-derived viruses in the total captured virome (SDs from mean = 2.2676, Fig. 3b, Table 2), suggesting that the majority of these viruses are not polyadenylated when captured in the host sequencing. Interestingly, we noticed a significant variation in the percentage of *A. salina*-derived viruses in polyA-selected libraries, with the mean of 14.06%, which could be a result of differences in the sample preparation prior to library generation, i.e., how long the animals were deprived of food before RNA extraction. Of note, we also detected fragments of four *Artemia*-derived viruses in the embryonic dataset, however, the very low overall yield of mapped reads suggests that these fragments might be parentally deposited products of viral sequence degradation.

**Table 2.** Summary of remapping results including the total number of reads mapped to assembled viral contigs from the merged dataset and number of reads mapped to putative *Artemia salina* viruses; SDs – number of standard deviations from mean calculated on the number of reads normalized to sequencing depth.

|  | Babonis<br>*et al.* | Tulin<br>*et al.* | Oren<br>*et al.* | Schwaiger<br>*et al. polyA* | Schwaiger<br>*et al. rRNA-*<br>*depleted* | Fidler<br>*et al.* | Warner<br>*et al.* |
| --- | --- | --- | --- | --- | --- | --- | --- |
| All reads<br>aligned to<br>viruses | 9,544 | 388 | 19,676 | 74,721 | 140,782 | 2,429 | 28,732 |
| SDs from<br>mean | -0.5133 | -0.5837 | -0.0156 | -0.1597 | 2.218 | -0.5154 | -0.4302 |
| Reads aligned<br>to *A. salina*<br>viruses | 1,362 | 63 | 721 | 1,783 | 127,633 | 1,116 | 537 |
| SDs from<br>mean | -0.3805 | -0.3908 | -0.3705 | -0.3818 | 2.2676 | -0.3547 | -0.3894 |

**Figure 3.**
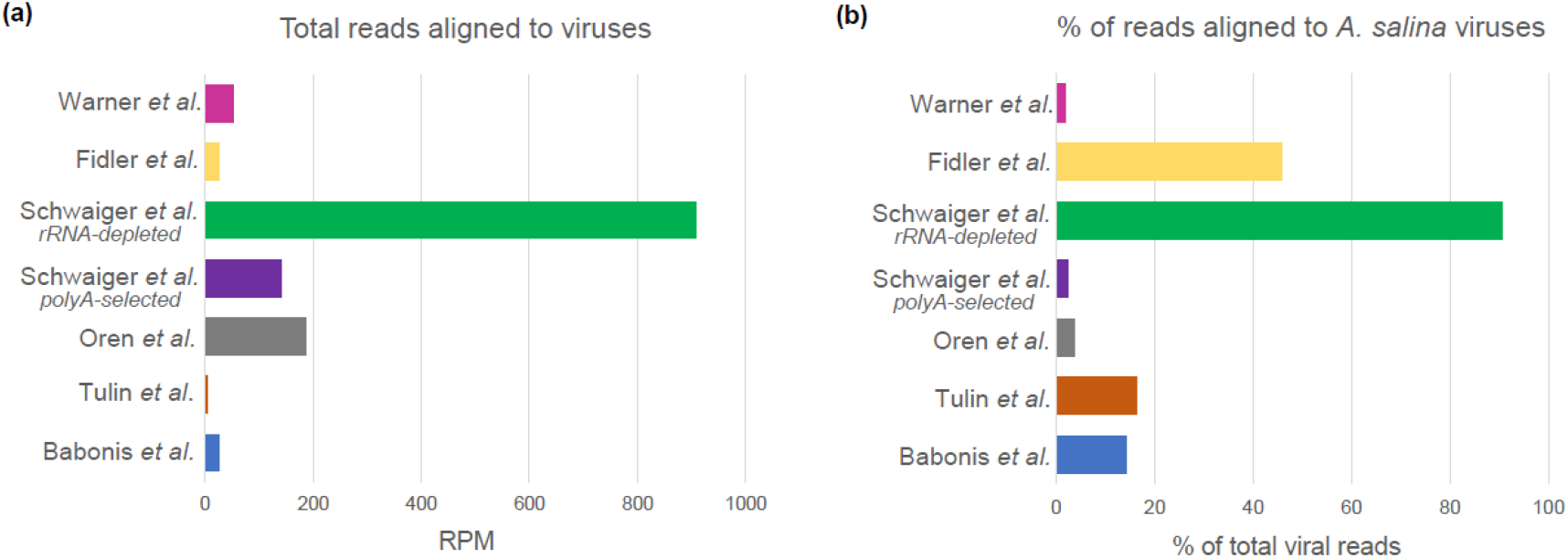
The number of reads mapped to *Nematostella* viral sequences assembled from all merged datasets presented as Reads Per Million (RPM) (a) and the fraction of total reads which mapped to *A. salina* viruses (b).

Comparison of viral communities in individual non-embryonic datasets suggested that while the overall composition of the virome displays a stable pattern across the samples, different lab populations might carry unique viruses, not found in other *Nematostella* groups. To test this assumption, we compared 2 datasets from adult animals for which 3 biological replicates were available (Babonis *et al.*, denoted as group 1 and Warner *et al.* denoted as group 7). PCA factor analysis revealed two factors with eigenvalue higher than 2 (3.161 and 2.278), which explained 52.69% and 37.97% of observed variance, respectively (Supplementary file 2). Statistical analysis of the extracted component loadings revealed a dual character of the observed variation between the two sets. The first factor showed no statistically significant differences and clustered these datasets together (t_2.854_ = 0, *P* = 1), while the analysis of the second factor suggested strong implicit variance between the studied groups (t_2.159_ = −12.3, *P* = 0.005). The result of partial overlap in viral sequences between two lab populations is not surprising in the light of the previously described *Nematostella* core virome. However, the strong variation we detected between the two datasets seems to confirm the significant contribution of the unique viruses in the studied lab populations.

## 4. Discussion

In the current study we identified 94 different sequences with sound homology to the known viruses from several viral families. Multiple-step removal of *N. vectesis*-mapping reads from the RNA-seq libraries resulted in the exclusion from the study of genome-integrated virus-derived sequences, such as retroviruses and endogenous-viral elements (EVEs). As no enrichment technique for viral particles, such as ultracentrifugation or size-based filtration, have been applied to any of the analyzed datasets, our search for fragments of exogenous viral genomes was relatively unbiased [4]. Nevertheless, it is important to note that the majority of analyzed datasets have been generated through mRNA enrichment by oligo(dT) selection, as these datasets were primarily designed for the whole transcriptome analysis. This, in turn, biased our search towards an increase in polyadenylated sequences present in some ssRNA(+) viruses [44], mRNA of RNA and DNA viruses [45, 46], viruses targeted by a host for degradation [47, 48], or products of other uncharacterized host-virus interface [49]. As expected, we have observed a significantly higher number of virus-mapping reads in the adult female library where rRNA was depleted with RiboMinusTM treatment (dataset “Schwaiger *et al.* rRNA-depleted”) when compared to the rest of the datasets. However, it needs to be taken into account that commercially available probes used for rRNA depletion are not designed for non-bilaterian animals and are less efficient in depleting the 5S small ribosomal subunit due to its high sequence variability between different animal phyla [50], which might compromise the library depth available to low-frequency viruses. Therefore, a truly unbiased search for RNA viruses would certainly gain from an rRNA depletion method custom-fitted for *N. vectensis* sequences.

Despite these limitations, we were able to retrieve a relatively broad representation of *N. vectensis* viruses, composed of 94 viral sequences, 21 of which were common to all non-embryonic datasets. The most common family present in almost all non-embryonic datasets was *Iridoviridae*, which belongs to the group of linear dsDNA viruses. Known hosts of *Iridoviridae* include amphibians, fish, reptiles, insects and crustaceans [51]. None of the identified sequences of *Iridoviridae* representatives was neither mapped in *Artemia* libraries, nor amplified from *Artemia* cDNA, which confirms that these dsDNA viruses are actively expressed and specific to *Nematostella*, or organisms comprising this holobiont, rather than food derived. Interestingly, members of *Iridoviridae* family were missing from all viromes available for other sea anemones species: *E. pallida* [16], *Actinia equina* [52] and *Bolocera sp* [4]. It seems plausible that such a major difference in the viral community composition between sea anemones might stem from the very distinct environmental conditions of these marine animals and therefore, a different ensemble of neighboring viral species. While *Bolocera* is an open-sea sea anemone [53], both *Actinia* and *Exaiptasia* occupy predominantly the intertidal zones which experience recurring but short-term fluctuations of water level and exposure to air [54]. In contrast, *Nematostella* is inhabiting mostly brackish lagoons of the east coast of North America, where it can be found burrowed into sand and mud [55]. Moreover, shallow waters of this habitat possess reduced buffering properties and expose *Nematostella* to strong shifts of environmental conditions throughout the year, as well as quite unique biota [56, 57], which altogether might result in the altered susceptibility of *N. vectensis* to different viral species.

An overall comparison between the four available viromes of the sea anemones revealed a general similarity of *N. vectensis* to *A. equina* and *Bolocera sp*., while the viral community of *E. pallida* displayed considerable differences. For instance, in the *A. equina* dataset we found two novel viruses which display the highest homology to viral sequences found in *N. vectensis* (Caledonia beadlet anemone dicistro-like virus 3 isolate B and A, homology to Wenzhou picorna-like virus 28 and Beihai picorna-like virus 71, respectively) [52]. The same number of viral sequences were common between our data dataset and *Bolocera sp*. virome (Beihai picorna-like virus 70 and Beihai picorna-like virus 118) [4]. In both cases, sequences identified in our study only partially covered the described viral genomes. Another similarity emerges from the distribution of viral families across studies. In the data from *A. equina* and *Bolocera* the most prevalent group are picorna-like viruses (50% and 59.1%, respectively), which falls into a novel Picorna-Calici clade established in Shi *et al.* [4]. Similarly, we identified in *Nematostella* dataset 37.2% viral sequences which belong to the Picorna-Calici clade, although the vast majority of them are classified here as “other viruses” due to the applied ICTV classification. On the contrary, *E. pallida* had more diverse viral community composition, in which among 40 identified viral families *Picornaviridae* constitute only 9.87% [16]. Moreover, we did not observe any viruses with obvious homology shared between *N. vectensis* and *E. pallida*. Finally, the most common family of *Exaiptasia* virome, *Herpesviridae*, was not found in any of the remaining sea anemones. Herpesviruses have been previously associated with other cnidarian species [3, 58, 59] and with *Symbiodinium microadriaticum* found in corals [14]. Given that *E. pallida* is also a host to several members of *Symbiodinium* family [60] and the contribution of *Herpesviridae* decreases in aposymbiotic state when compared to a fully symbiotic *Exaiptasia* (8.1% and 12.9%, respectively) [16], it is possible that this viral family is associated with the presence of these symbionts and hence, not found in sea anemone species which do not harbor zooxanthellae.

Among 21 viral sequences present in all non-embryonic datasets which we denominated as the core virome of *Nematostella*, 6 sequences homologous to 4 different viruses were identified in its primary source of food, *A. salina* nauplii. Unsurprisingly, known hosts of those viruses include insects, crustaceans as well as insect and vertebrate parasitic nematodes [4]. Interestingly, we were able to amplify by RT-PCR fragments of two RNA viruses included in our core genome (Sanxia water strider virus 10 and Hubei sobemo-like virus 41) from the cDNA of *A. salina*, while not detecting any matching reads in none of the two replicates of *Artemia* RNA-seq libraries. The most plausible explanation is that the lack of polyA tail on the 3’ end of the RNA molecule would hinder their detection in the transcriptome analysis but would not bias a cDNA preparation constructed with random hexamers. Therefore, we cannot exclude the fact that the overlap between food-derived viruses and *Nematostella* holobiont-specific virome maybe be more significant than described here and a less biased sequencing approach is needed to fully characterize it. However, the fact that the majority of viral sequences targeted by RT-PCR were amplified from adult *Nematostella* but not from *Artemia nauplii* (Figure S1) is a strong indication that many of the viruses we detected in the RNA-seq are not food-derived. The presence of persistent or prevalent viruses in lab populations of model animals was shown before for *Drosophila* [61] and very recently was also reported for zebrafish [62].

Besides the presence of a stable core virome of *Nematostella*, we have detected several population-specific viruses. Namely, 5 out of 7 analyzed datasets possess unique fragments of viral genomes, not found in any other dataset. Such specificity was previously reported on a species level within the cnidarian genus of *Hydra* [3], although the species-specific diversity was associated with an extensive ensemble of bacteriophages, which were not the subject of our study. In the case of two of the datasets analyzed here, the contribution of population-specific viruses was remarkable and reached 52.5 – 93.03% of the total reads mapping to viruses (datasets “Babonis *et al.*” and “Oren *et al.*”, respectively). Interestingly, the unique virus detected in the tissue-specific dataset (“Babonis *et al.*”) exhibits the highest homology to a virus identified previously in tunicates [4] and it is unevenly clustering in only one library replicate generated from the mesentery tissue. In natural populations, such a virome diversity could reflect unique environmental conditions. However, to the best of our knowledge, there are no significant differences in *N. vectensis* culture between different lab populations as they originate from the same population from Rhode River, MD, USA, which was cultured and used for the genome sequencing [38]. All together, we cannot exclude the possibility that this unique virus might not represent a stable population-unique viral community, but instead, it was acquired from other species cultured in the research facility where the *Nematostella* polyps were kept.

Finally, our analysis of Tulin *et al.* dataset which spans 24 hours of embryonic development revealed that the viral load in this early life stage library was negligible. None of the sequences from the core virome specific to *N. vectensis* was present neither in the Tulin *et al.* RNA-seq dataset [30], nor in our early planulae cDNA preparation. Comparison to the individual assemblages of available early developmental stages, i.e., from an unfertilized egg, blastula and gastrula confirmed this pattern suggesting that the lab *Nematostella* is free of viruses in its early developmental stage and acquires them throughout life, both by food ingestion and uncharacterized ways of entry. Unfortunately, the viral datasets of other sea anemones do not span multiple developmental stages which impedes a direct comparison of their embryonic viral load. Unexpectedly, the only viral sequences assembled from the data by Tulin *et al.* 2013, which are similar to the Rous sarcoma virus [63], exhibited remarkable homology (99% identity at the nucleotide level) to transcriptomic sequences from the reef-building coral *Acropora millepora* [64] and the stalked jellyfish *Haliclystus sanjuanensis*. Such an unusually high level of homology among species that separated more than 600 million years ago [65] raises the possibility of contamination. Of note, the homology level of the *N. vectensis* sequences to the Rous sarcoma virus and other closely-related vertebrate viruses (e.g. Avian leukosis virus) was lower (<97%) than the homology among these three far-related cnidarians. However, as these sequences failed to be filtered out by multiple steps of mapping to *Nematostella* genome and were missing from individual early-developmental assemblages, this strengthens our prediction that they might represent a contamination of the RNA-seq libraries rather than a genome-integrated cnidarian retrovirus.

Although most studies are focusing on the role of viruses in the pathogenesis of vertebrates, there is an increasing understanding of the significance of viral communities in the formation of a stable holobiont among all living organisms. Here, we re-analyzed several high-throughput RNA-seq datasets available for a cnidarian model organism, *Nematostella vectensis* and we developed a straightforward approach of *de novo* assembly followed by a multi-step, homology-based virus identification. Our study revealed a diverse set of eukaryotic, non-integrated viruses spread across 7 different lab populations, among which we identified both the core virome present in all datasets and several population-unique viruses. The observed absence of viral community during the early stages of development and identification of viruses shared with the primary food source of *N. vectensis*, *A. salina*, provides an initial insight into the course of viral community acquisition in *Nematostella vectensis*. Further research combining both non-targeted and virus-enriched deep sequencing approach is essential for a full characterization of *Nematostella* viral community.

## Supplementary Materials

The following are available online, Supplementary file 1: Table S1: Detailed list of NCBI accession numbers, raw and retained reads counts. Table S2: List of contigs used for RT-PCR validation of *Nematostella vectensis* common virome. Table S3: Identified viral sequences in the merged dataset composed of all RNA-seq libraries pooled together. Table S4: Identified viral sequences in the two replicates of *A. salina* RNA-seq libraries. Table S5: Core virome of all the non-embryonic datasets. Table S6: Identified viral sequences in the Babonis *et al.* dataset. Table S7: Identified viral sequences in the Tulin *et al.* dataset. Table S8. Identified viral sequences in the Oren *et al.* dataset. Table S9. Identified viral sequences in the joined dataset from Schwaiger *et al.* and Vienna University submission, libraries generated by polyA selection. Table S10. Identified viral sequences in the joined dataset from Schwaiger *et al.* and Vienna University submission, libraries generated by rRNA depletion. Table S11: Identified viral sequences in the Fidler *et al.* dataset. Table S12: Identified viral sequences in the Warner *et al.* dataset. Text file S1: Viral sequences in the merged dataset, trimmed. Text file S2: Viral sequences in the merged dataset, untrimmed. Text file S3: Viral sequences in the Babonis *et al.* dataset, trimmed. Text file S4: Viral sequences in the Babonis *et al.* dataset, untrimmed. Text file S5: Viral sequences in the Tulin *et al.* dataset, trimmed. Text file S6: Viral sequences in the Tulin *et al.* dataset, untrimmed. Text file S7: Viral sequences in the Oren *et al.* dataset, trimmed. Text file S8: Viral sequences in the Oren *et al.* dataset, untrimmed. Text file S9: Viral sequences in the Schwaiger *et al.* and Vienna University submission dataset, polyA selected, trimmed. Text file S10: Viral sequences in the Schwaiger *et al.* and Vienna University submission dataset, polyA selected, untrimmed. Text file S11: Viral sequences in the Schwaiger *et al.* and Vienna University submission dataset, rRNA depleted, trimmed. Text file S12: Viral sequences in the Schwaiger *et al.* and Vienna University submission dataset, rRNA depleted, untrimmed. Text file S13: Viral sequences in the Fidler *et al.* dataset, trimmed. Text file S14: Viral sequences in the Fidler *et al.* dataset, untrimmed. Text file S15: Viral sequences in the Warner *et al.* dataset, trimmed. Text file S16: Viral sequences in the Warner *et al.* dataset, untrimmed. Supplementary file 2: PCA analysis. Figure S1: Validation of presence of candidate viruses by RT-PCR.

## Author Contributions

conceptualization, M.L. and Y.M.; methodology, M.L..; software, Y.H..; validation, M.L.; formal analysis, M.L.; investigation, M.L.; resources, Y.M.; data curation, M.L.; writing—original draft preparation, M.L..; writing—review and editing, Y.H. and Y.M..; visualization, M.L.; supervision, Y.M.; project administration, Y.M.; funding acquisition, Y.M.

## Acknowledgments

We would like to thank Dr. Shelby Rinehart (Hebrew University of Jerusalem) for performing the statistical analysis and Dr. Vengamanaidu Modepalli (Marine Biological Association of UK) for the help in the design of the bioinformatic pipeline. We are also grateful to Dr. Michal Bronstein and Ms. Adi Turjeman (The Center for Genomic Technologies, Hebrew University of Jerusalem) for their help with transcriptome sequencing.

