## Supplementary material for "Initial virome characterization of the common cnidarian lab model *Nematostella vectensis*": Figure S1

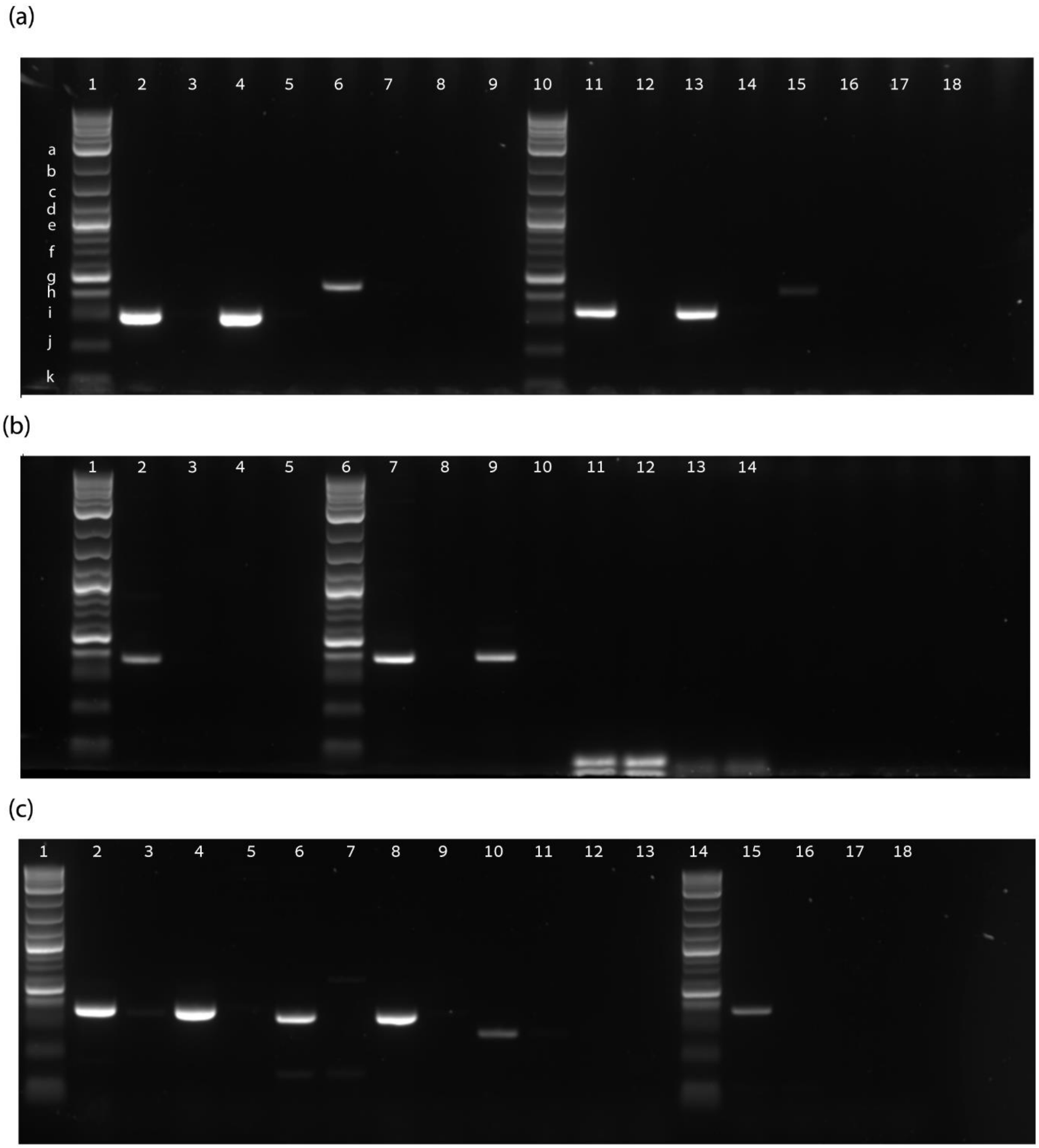


**Figure S1.** Validation of presence of candidate viruses by RT-PCR. **(a)** lanes 2,6,11,15 – adult *Nematostella*, 3,7,12,16 – planula *Nematostella*, 4,8,13,17 – *A. salina*, 5,9,14,18 – negative controls; lanes 2-5 – c40691_g1_i1, 6-9 – c145931_g1_i1, 11-14 – c99447_g1_i1, 15-18 – c44518_g1_i1. **(b)** lanes 2,7,11 – adult *Nematostella*, 3,8,12 – planula *Nematostella*, 4,9,13 - *A. salina*, 5,10,14 – negative controls; lanes 2-5 – c18556_g1_i1 , 7-10 – c101721_g1_i1, 11-14 - NVE5273; **(c)** lanes 2,6,10,15 - adult *Nematostella*, 3,7,11,16 - planula *Nematostella*, 4,8,12,17 - *A. salina*, 5,9,13,18 – negative controls; lanes 2-5 – c100306_g1_i1, 6-9 – c102334_g1_i1, 10-13 – c101934_g2_i1, 15-18 – c97783_g2_i1. Ladder legend: **a** – 3 kb, **b** – 2 kb, **c**- 1.5 kb, **d** – 1.2 kb, **e** – 1 kb, **f** – 0.7 kb, **g** – 0.5 kb, **h** – 0.4 kb, **i** – 0.3 kb, **j** – 0.2 kb, **k** – 0.1 kb.
