## Supplementary file 2 for "Initial virome characterization of the common cnidarian lab model *Nematostella vectensis*"

### ▼File: Untitled1.syz

IMPORT successfully completed.

### ▼Factor Analysis

#### Latent Roots (Eigenvalues)

| 1 | 2 | 3 | 4 | 5 | 6 |
|---|---|---|---|---|---|
| 3.161 | 2.278 | 0.389 | 0.151 | 0.014 | 0.006 |

#### Component Loadings

| | 1 | 2 | 3 | 4 | 5 |
|---|---|---|---|---|---|
| **C2** | 0.754 | -0.635 | 0.005 | 0.156 | -0.029 |
| **C3** | 0.820 | -0.444 | 0.199 | -0.301 | 0.000 |
| **C4** | 0.592 | -0.774 | -0.189 | 0.110 | 0.029 |
| **C5** | 0.659 | 0.561 | -0.491 | -0.097 | 0.009 |
| **C6** | 0.764 | 0.625 | 0.115 | 0.071 | -0.078 |
| **C7** | 0.743 | 0.610 | 0.243 | 0.099 | 0.079 |

#### Variance Explained by Components

| 1 | 2 | 3 | 4 | 5 |
|---|---|---|---|---|
| 3.161 | 2.278 | 0.389 | 0.151 | 0.014 |

#### Percent of Total Variance Explained

| 1 | 2 | 3 | 4 | 5 |
|---|---|---|---|---|
| 52.690 | 37.968 | 6.484 | 2.522 | 0.235 |

Scree Plot

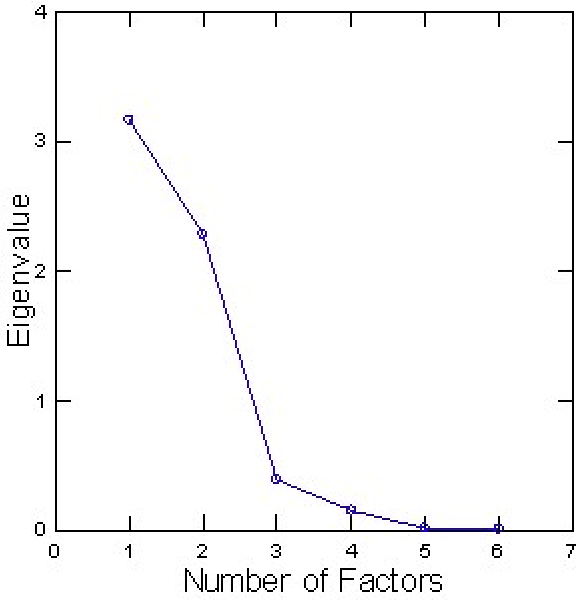

### Factor Loadings Plot

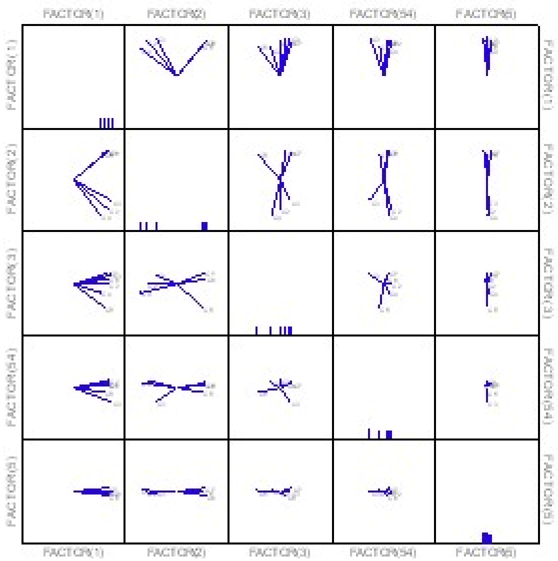

▶ Factor Analysis

▼ Hypothesis Testing: Two-sample t-test

**Two-sample t-test on PCA1 Grouped by PROJECT vs Alternative = 'not equal'**

| GROUP | N | Mean | Standard Deviation |
|---|---|---|---|
| 1 | 3 | 0.722 | 0.117 |
| 7 | 3 | 0.722 | 0.056 |

Separate Variance

| | | |
|---|---|---|
| Difference in Means | : | 0.000 |
| 95.00% Confidence Interval | : | -0.246 to 0.246 |
| t | : | 0.000 |
| df | : | 2.854 |
| p-value | : | 1.000 |

Pooled Variance

| | | |
|---|---|---|
| Difference in Means | : | 0.000 |
| 95.00% Confidence Interval | : | -0.208 to 0.208 |
| t | : | 0.000 |
| df | : | 4.000 |
| p-value | : | 1.000 |

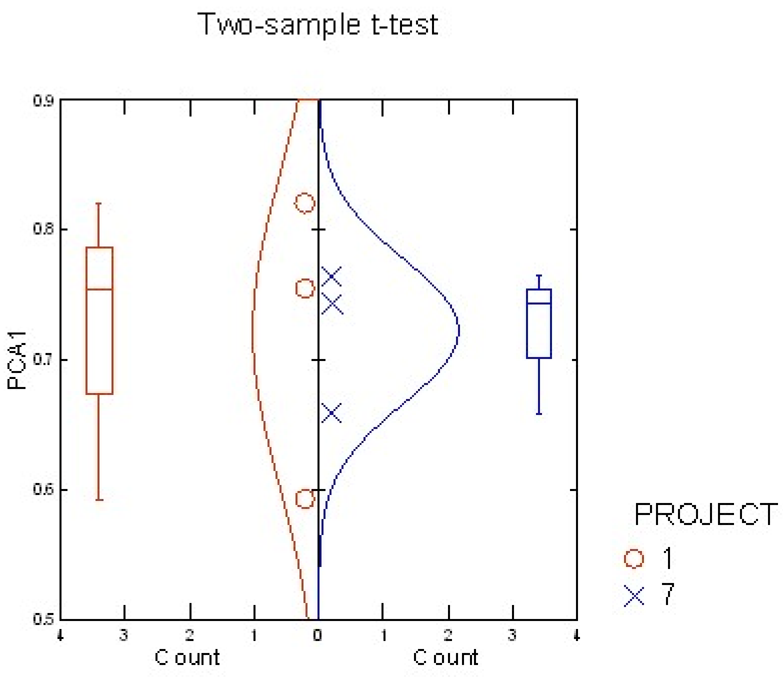


▼ Hypothesis Testing: Two-sample t-test

**Two-sample t-test on PCA2 Grouped by PROJECT vs Alternative = 'not equal'**

| GROUP | N | Mean | Standard Deviation |
|---|---|---|---|
| **1** | 3 | -0.616 | 0.168 |
| **7** | 3 | 0.599 | 0.033 |

Separate Variance

| | | |
|---|---|---|
| Difference in Means | : | -1.215 |
| 95.00% Confidence Interval | : | -1.611 to -0.819 |
| t | : | -12.300 |
| df | : | 2.159 |
| p-value | : | 0.005 |

Pooled Variance

| | | |
|---|---|---|
| Difference in Means | : | -1.215 |
| 95.00% Confidence Interval | : | -1.489 to -0.941 |
| t | : | -12.300 |
| df | : | 4.000 |
| p-value | : | 0.000 |

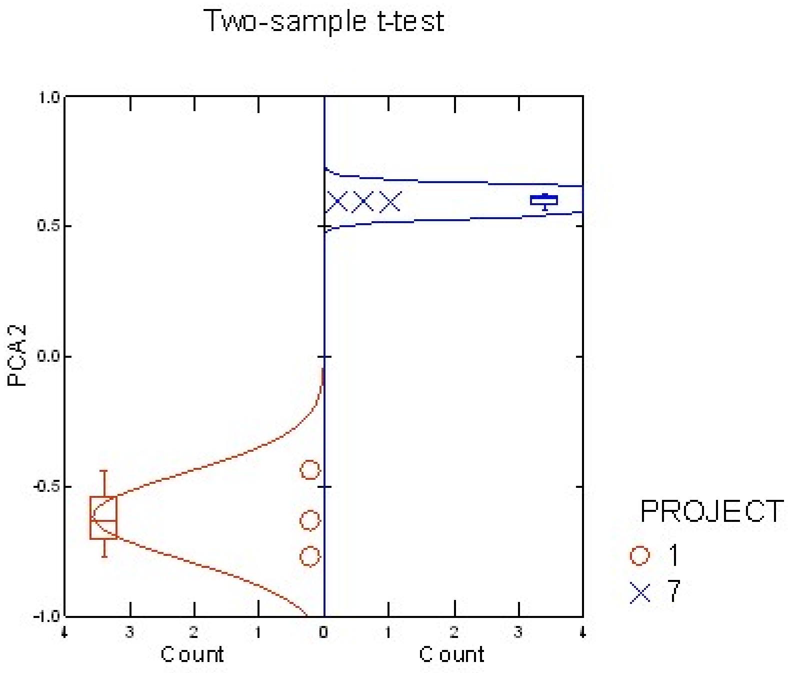
